# Impaired hippocampal representation of place in the *Fmr1*-knockout mouse model of Fragile X syndrome

**DOI:** 10.1101/191775

**Authors:** Tara Arbab, Cyriel MA Pennartz, Francesco P Battaglia

**Author notes:** **Corresponding Author** Tara Arbab.

## Abstract

Fragile X syndrome (FXS) is an X-chromosome linked intellectual disability and the most common genetic cause of autism spectrum disorder (ASD). Building upon demonstrated deficits in neuronal plasticity and spatial memory in FXS, we investigated how spatial information processing is affected *in vivo* in an FXS mouse model (*Fmr1*-KO). Healthy hippocampal neurons (so-called place cells) exhibit place-related activity during spatial exploration, and the stability of these spatial representations can be taken as an index of memory function. We find impaired stability and reduced specificity of *Fmr1*-KO spatial representations. This is a potential biomarker for the cognitive dysfunction observed in FXS, informative on the ability to integrate sensory information into an abstract representation and successfully retain this conceptual memory. Our results provide key insight into the biological mechanisms underlying cognitive disabilities in FXS and ASD, paving the way for a targeted approach to remedy these.

## Introduction

The most common genetic cause of autism spectrum disorder (ASD) is Fragile X syndrome (FXS) ^1,2^: an intellectual disability in which expression of the fragile X mental retardation protein (FMRP) is silenced, resulting in disturbed neuronal communication ^3,4^. Due to its simple genetic etiology, FXS shows promise for understanding neuropsychiatric disease from genes, to circuits, to cognitive impairment ^5^. Particularly affected in FXS human patients and animal models is the hippocampus ^6,7^, a brain structure essential for consolidating experiences into conceptual and spatial memory ^8-11^. When healthy humans and animals explore a space, hippocampal neurons (so-called place cells) exhibit place-related activity ^12,13^. The stability of these spatial representations can be taken as an index of memory function ^14,15^. It follows that anomalous place cell activity in disease models may be characteristic of cognitive impairment in neurological disorders. We used a spatial exploration paradigm to investigate in an FXS mouse model (*Fmr1*-KO) ^16,17^ how spatial information processing is affected *in vivo* by recording hippocampal place cell activity.

## Results and Discussion

### Fmr1-KO does not affect exploratory behavior

We recorded neuronal activity in hippocampal CA1 (Figure 1A) during four subsequent spatial exploration sessions (across two days) in five *Fmr1*-KO mice and five WT control mice (Figure 1B). The first two recording sessions (on the morning and afternoon of the first day) and the third recording sessions (on the morning of the second day) were done with a complete set of four visual cues marking the environment (“Full cue” sessions). For the fourth session (on the afternoon of the second day), three of these visual cues were removed from the room, leaving an incomplete set of cues by which the animal could localize itself (“Probe” session). Our recordings yielded 122 WT and 141 *Fmr1*-KO putative pyramidal cells.

**Figure 1.**
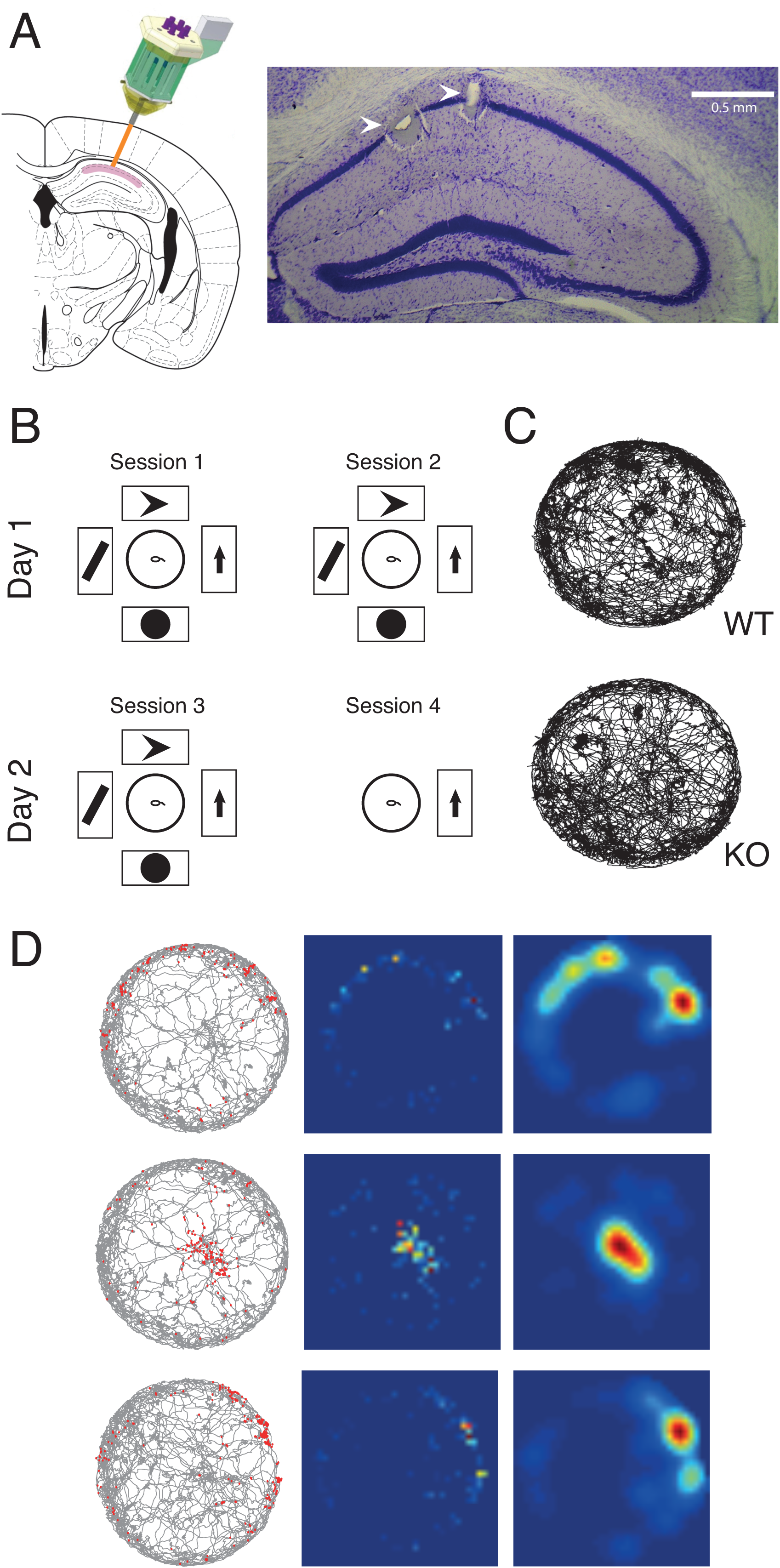
Experimental Setup. (A) Histology. Left, schematic of microdrive implantation target. Right, coronal section showing the recording locations of two tetrodes in dorsal hippocampus CA1 (arrowheads). (B) Schematic of the behavioral protocol. On two consecutive days, in two sessions per day, animals freely explored a circular open field arena (middle). The arena was surrounded by four posters of geometric figures in the first three sessions, and only one poster in the fourth session. (C) Behavior. Accumulated trajectories of a WT and a KO animal exploring the arena during an example session. (D) Three example firing rate maps. Left, accumulated trajectory of animal exploration during a session, with spikes recorded from a single pyramidal cell superimposed in red. Middle, heat map of these spikes created by binning and normalizing this data. Right, smoothed heat map of these binned and normalized spikes.

Hippocampal place fields are thought to result from a complex process of integration of different types of information during exploration of an environment. Instantaneous multisensory inputs from the environment are combined into a ‘scaffold’ of internally generated representations, which depend to a large extent on self-motion information ^18,19^. Therefore, we first controlled for parameters of exploratory behavior, which may act as a confounding factor for neuronal processing of spatial information (Figure 1C). We found no difference between WT and *Fmr1*-KO running speed (WT mean=5.99cm/s, SEM=0.16; KO mean=5.48cm/s, SEM=0.20, P=0.15) and thigmotaxis (average distance from the center of the arena, WT mean=25.29cm, SEM=0.31; KO mean=24.36cm, SEM=0.49, P=0.10) across sessions. Additionally, there was no difference in the maximum and mean pyramidal cell firing rate across genotypes (Table S1).

### Fmr1-KO reduces spatial specificity of place cells

Pyramidal cells of both genotypes exhibited spatially selective activity: place fields (Figure 1D). We found no difference between WT and *Fmr1*-KO mice in the number of place fields per cell, or the spatial information that each spike carried. However, *Fmr1*-KO place fields were significantly larger than those of wildtype animals, based both on counting active pixels in the normalized map (WT mean=9429, SEM=18; KO mean=9549, SEM=16, P<0.001) and by comparing the place fields in the smoothed maps (Table S1).

We determined spatial specificity of place cells as the firing rate increase of each cell within its field (Figure 2). Spatial specificity of *Fmr1*-KO pyramidal cells was significantly reduced compared with WT (Bonferroni-corrected two-way genotype x session ANOVA, effect of genotype: F_1,586_ = 25.55; P < 0.0001). There was no effect of session and no interaction effect across cells.

**Figure 2.**
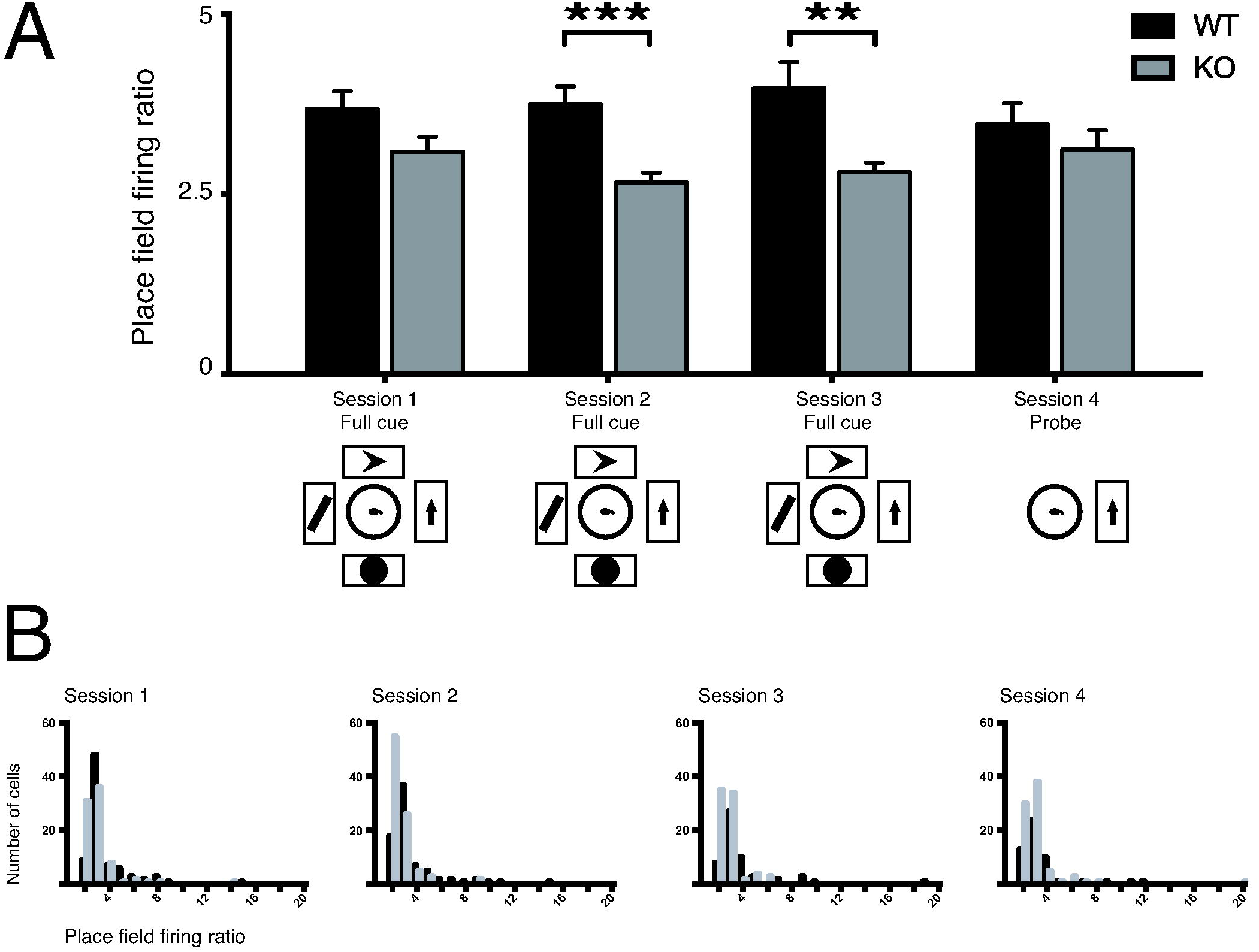
Spatial specificity of place cells per session. (A) Place field firing rate increase of WT (black) and *Fmr1*KO (gray) pyramidal cells. Data are represented as mean [INLINE] SEM. Session 1: WT 80 cells, mean=3.69, SEM=0.22; KO 81 cells, mean=3.09, SEM=0.18. Session 2: WT 77 cells, mean=3.75, SEM=0.25; KO 91 cells, mean=2.66, SEM=0.12. Session 3: WT 55 cells, mean=3.97, SEM=0.38; KO 78 cells, mean=2.81, SEM=0.10. Session 4: WT 52 cells, mean=3.47, SEM=0.28; KO 80 cells, mean=3.12, SEM=0.24. (B) Distributions of place field firing rate increase of WT (black) and *Fmr1*-KO (gray) pyramidal cells. ** P < 0.01; *** P < 0.001

### Fmr1-KO impairs short-term stability of spatial representation

To assess the stability of place cell activity, we first evaluated the similarity of the firing rate maps between the halves of each session (Figure 3). Pixel-based Pearson correlations between rate maps were significantly reduced in *Fmr1*-KO mice compared with WT (Bonferroni-corrected two-way genotype x session ANOVA, effect of genotype: F_1,590_ = 46.34; P < 0.0001). We found no effect of session and no interaction effect. These results were consistent when data were analyzed by comparing per-animal means (Bonferroni-corrected two-way genotype x session ANOVA, effect of genotype: F_1,30_ = 5.27; P = 0.03). Comparison between the quarters of each session yielded the same conclusion: correlations between rate maps were significantly reduced in *Fmr1*-KO mice compared with WT (Bonferroni-corrected two-way genotype x session ANOVA, effect of genotype: F_1,1457_ = 92.58; P < 0.0001) with no effect of session and no interaction effect (Table S2).

**Figure 3.**
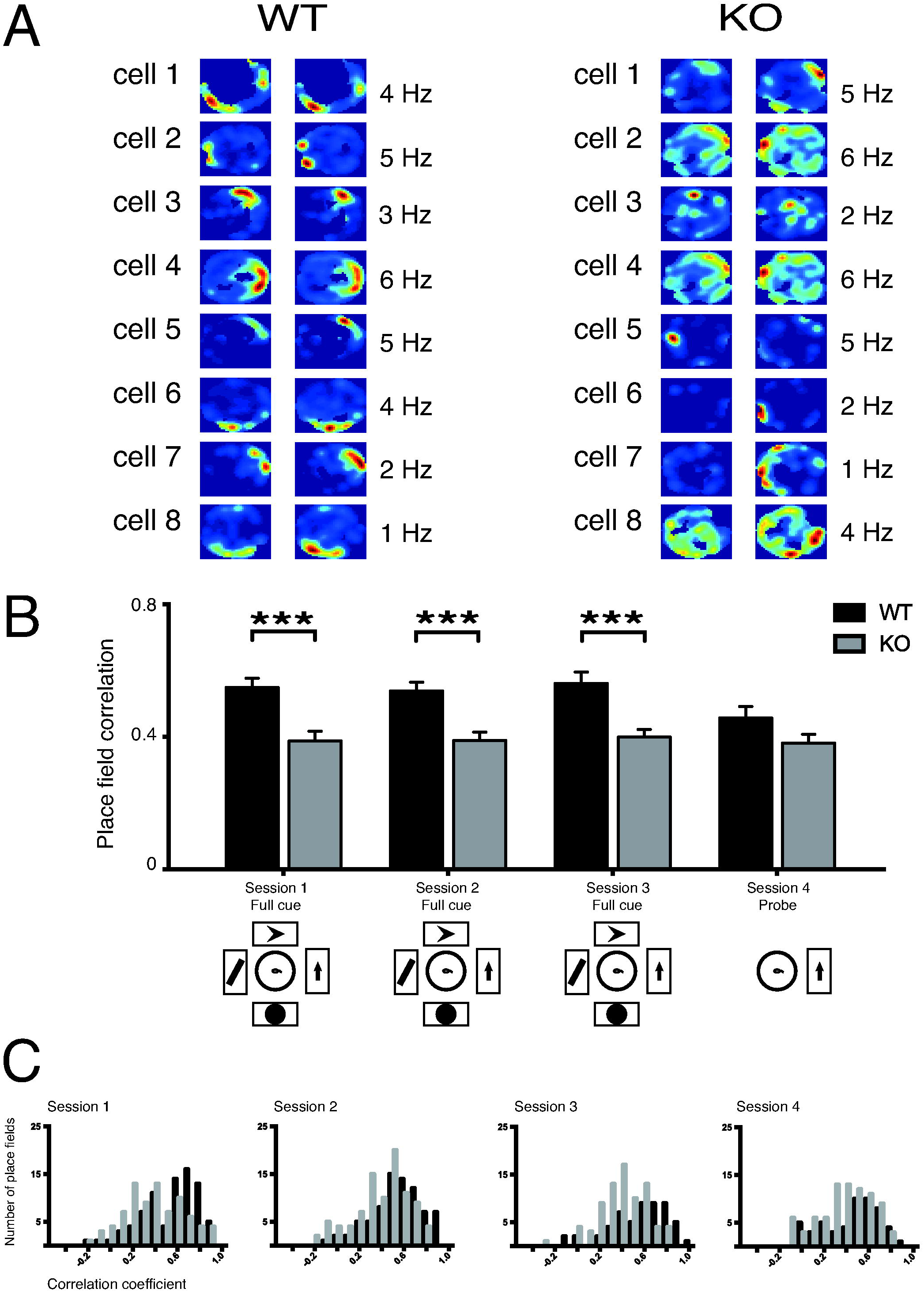
Stability of firing rate maps within sessions. (A)Example WT and KO firing rate maps (taken from all sessions), split between the first and second halves of each recording session (left and right panels), to illustrate the stability of each map. Each heat map is scaled by the maximum firing rate (indicated in Hz) of the cell within that session. (B) Correlation of WT (black) and *Fmr1*-KO (gray) firing rate maps. Data are represented as mean [INLINE] SEM. Session 1: WT 80 fields, mean=0.55, SEM=0.028; KO 81 fields, mean=0.39, SEM=0.03. Session 2: WT 78 fields, mean=0.54, SEM=0.03; KO 91 fields, mean=0.39, SEM=0.03. Session 3: WT 55 fields, mean=0.56, SEM=0.03; KO 78 fields, mean=0.40, SEM=0.02. Session 4: WT 53 fields, mean=0.46, SEM=0.03; KO 82 fields, mean=0.38, SEM=0.03. (C) Distributions of the stability of WT (black) and *Fmr1*-KO (gray) pyramidal cells. *** P < 0.001.

Thus, the effects of the FXS mutation on spatial representation in *Fmr1*-KO mice are not attributable to behavioral differences or basic physiological properties such as pyramidal cell firing rate. However, the relative strength of firing of WT place cells within fields is greater than in *Fmr1*-KO mice, the latter shows increased size of place fields, and the location of place responses is less stable in *Fmr1*-KO mice than in WT controls in short intervals within recording sessions in the same environment.

As for any neural integration operation, self-localization is affected by the accumulation of errors, resulting in drift which increases with time ^20,21^. Spatially informative sensory cues can realign the drifting map, therefore reducing error. In interpreting our current findings, one possibility is that the FXS mutation affects the sensory information-dependent updating of the self-motion based map, while leaving the map itself relatively spared.

As sensory cues may rapidly induce profound changes in the spatial map ^22^, in an attention modulated way ^15^, the increased instability we observe in *Fmr1*-KO mice may be due to a lower weight of sensory inputs in determining place cell firing. Indeed, FXS patients show defective attention and integration of new information ^23^.

### Fmr1-KO spatial representation does not reflect changes in environment

To determine whether *Fmr1*-KO mice are impaired in sensory information-dependent updating of their spatial representation, we assessed the stability of firing rate maps between “Full cue” and “Probe” sessions on the same day. Here, we found no direct effect of genotype or session, but we found a significant interaction effect (Bonferroni-corrected two-way genotype x session ANOVA, interaction effect: F_1,281_ = 6.074; P = 0.0143). Specifically, *Fmr1*-KO pyramidal cells showed a significantly reduced correlation of activity compared with WT between the “Full cue” sessions (i.e., sessions 1 and 2, in which the open field arena was surrounded by the same four visual cues). The firing rate map correlation between the “Full cue” and the novel “Probe” (i.e., sessions 3 and 4, in between which three of these cues were removed) environment however, did not differ between genotypes (Figure 4).

**Figure 4.**
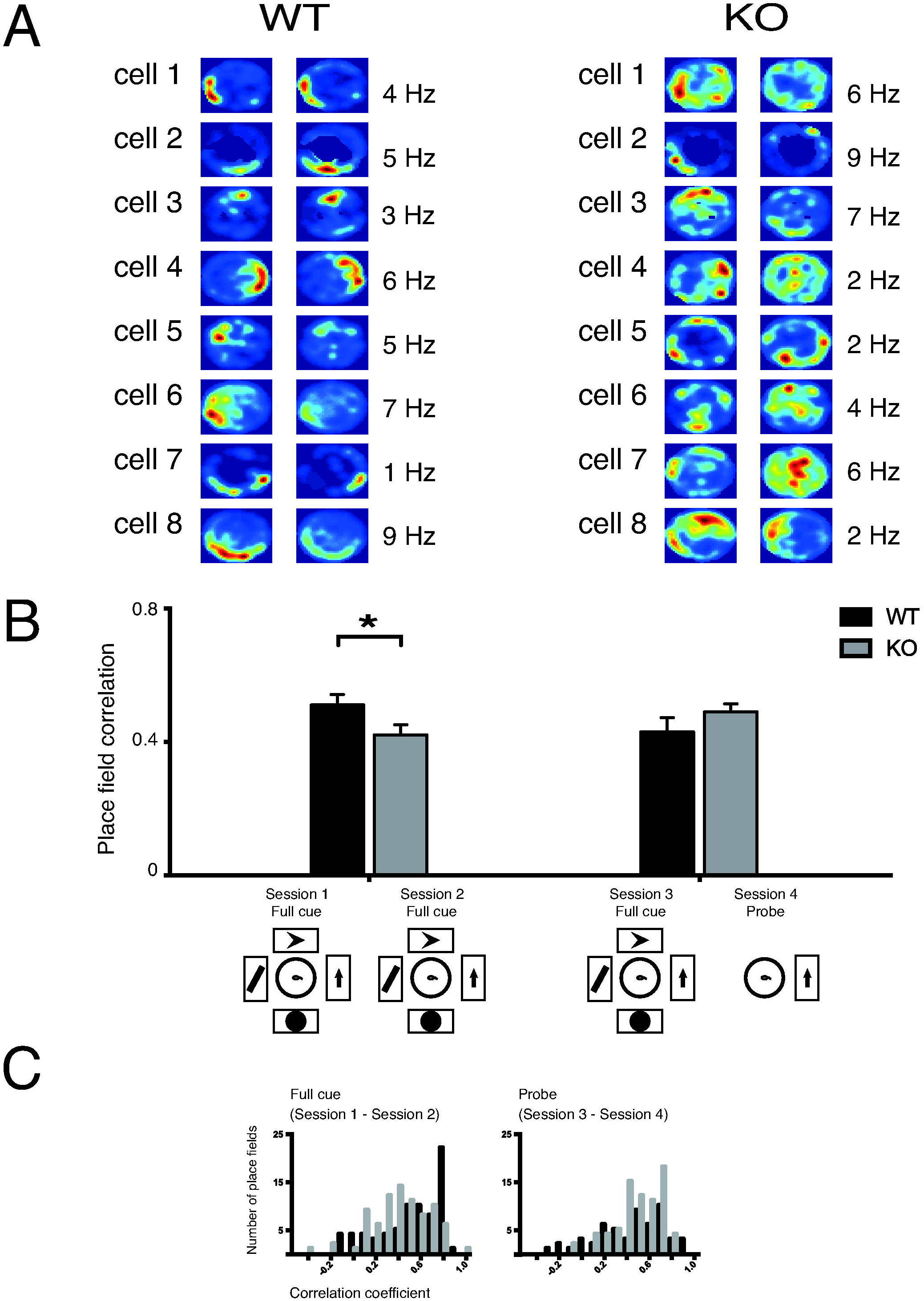
Stability of firing rate maps between sessions. (A)Example WT and KO firing rate maps (taken from all sessions), split between the two daily recording sessions (left and right panels), to illustrate the stability of each field. Each heat map is scaled by the maximum firing rate (indicated in Hz) of the cell within that session. (B) Correlation of WT (black) and *Fmr1*-KO (gray) firing rate maps. Data are represented as mean ± SEM. Full cue session: WT 75 fields, mean=0.51, SEM=0.03; KO 81 fields, mean=0.42, SEM=0.03. Probe session: WT 54 fields, mean=0.43, SEM=0.04; KO 75 fields, mean=0.49, SEM=0.02. (C) Distributions of the stability of WT (black) and *Fmr1*-KO (gray) pyramidal cells* P < 0.05.

This lack of difference between genotypes under the “Probe” condition corroborates the idea that WT place cell activity is more strongly affected by the changes in the environment, whereas *Fmr1*-KO place cells may preferentially rely on other (e.g. self-motion or internal ^24^) sources of information. This lack of difference in the probe condition cannot be ascribed to overnight memory consolidation effects because *Fmr1*-KO place cells were still impaired in session 3, which also took place the day after sessions 1 and 2.

### Spatial representation impairments in Fmr1-KO provide biomarker for FXS deficits

Both animal ^8,9^ and human ^10,11^ studies link the hippocampus to spatial, contextual, and autobiographical memory. In the same way that place cells in animals exploring an environment can encode that space, the activity of hippocampal neurons in humans can encode abstract representations of multi-sensory perceptual information ^13,25^. Hippocampal dysfunction is a critical component of intellectual pathologies such as FXS and ASD, in which impairments of conceptualization and memory are observed ^6,7,26^. Here, the delicate system that allows the brain to carefully fine-tune which information it retains is disrupted, because of the devastating effect on activity-dependent synaptic plasticity that underlies learning and memory. This ultimately contributes to anomalous processing of social and environmental cues and associated deficits in memory and cognition ^27^. Although they are equally affected neurologically, it has been difficult to assess these cognitive deficits in animal models with the same robustness as in human FXS patients ^28^. Our findings take the middle ground by demonstrating on a network level that altered physiology in *Fmr1*-KO leads to impaired hippocampal information processing.

There is a wide array of FXS physiological deficits which might underly our results. Stability of spatial representation requires long-term potentiation (LTP) associated with NMDA receptor activity in hippocampal CA1 ^29-32^. FMRP regulates subunit composition of hippocampal NMDA receptors ^33^ and may therefore contribute to *Fmr1*-KO pathophysiology by affecting synaptic plasticity through altered subunit composition of NMDA receptors. Indeed, LTP deficits are observed in *Fmr1*-KO mice ^34-37^. Additionally, *Fmr1*-KO mice show higher dendritic expression of the *HCN1* gene in hippocampal CA1 ^38^, which impairs spatial memory and plasticity in pyramidal neurons by affecting the ability of the entorhinal cortex to excite them ^39^ The instability we find in *Fmr1*-KO firing rate maps may be interpreted within this context as an increase in *HCN1*-mediated control over CA1 pyramidal cell plasticity from entorhinal inputs through the perforant pathway, which affects the sensory information-dependent updating of the self-motion based map as described above. Finally, disrupted network mechanisms ^40^ regulating the inflow of information between the hippocampus and entorhinal cortex ^41,42^ may contribute to improper routing of sensory information to the hippocampus, or in the failure to elicit spike-timing dependent plasticity ^32,43,44^. While the cognitive effects of these deficits have proven difficult to assess behaviorally in *Fmr1*-KO ^28^, we find that they may contribute to disrupting neural mechanisms that establish associations between external cues and internally generated or self-motion dependent representations.

Hippocampal place cells are one of the best understood systems in the brain where we have reached an initial understanding of the relationship between neural dynamics, information encoding, and cognition. Here we have shown that they may provide a powerful tool in understanding intellectual disability and ASD in a mouse model of FXS, in which it has been surprisingly difficult to demonstrate consistent cognitive deficits despite its clear genetic etiology. We find impaired specificity and stability of CA1 place cell activity in *Fmr1*-KO mice, both within and across subsequent exploration sessions, while these mice show a relatively spared place field response and their behavior and firing-rate parameters do not significantly differ from WT mice. Our results link impaired physiology with cognition more deeply than possible with traditional behavioral of physiological assays, and offer a potential biomarker for testing of therapeutic strategies.

## Methods

### Subjects

We used five *Fmr1*-KO mice ^17^ and five littermate wildtype (WT) control mice. All experiments were performed in accordance with Dutch National Animal Experiments regulations, were approved by the Universiteit van Amsterdam, and were carried out by certified personnel. Animals were received from the Erasmus Medisch Centrum Rotterdam breeding unit at an age of 8 weeks and group-housed until surgery. They were maintained on a regular 12-hour light-dark cycle (lights on: 8am, lights off: 8pm) and received standard food pellets and water *ad libitum* throughout the experiment. To minimize bias due to possible undetected changes in environmental conditions, *Fmr1*-KO and WT animals were always studied in pairs; both recordings were done on the same day and counterbalanced per genotype. Once habituated to the experimenter and handling, the mice underwent drive implantation surgery under buprenorphine-isoflurane anesthesia and were left to recover fully before the start of the experiment.

### Electrophysiological techniques

Six independently moveable tetrodes were loaded into a custom-made microdrive^32,45^ and implanted over the dorsal hippocampus (AP: -2.0mm, ML: -2mm; Figure 1A). The tetrodes were lowered into the CA1 pyramidal cell layer guided by electrophysiological signals (sharp wave-ripple events) over the course of days following implantation surgery. Electrophysiological activity was recorded on a 27-channel analog Neuralynx data acquisition system at a 32kHz sampling rate. Tetrode signals (bandpass filtered 0.6-6.0kHz) were referred to a nearby tetrode which was targeted to a location devoid of single unit activity. Single-unit data were preprocessed with Klustakwik ^46^ for automated spike clustering and the results were manually refined using Klusters ^47^. The resulting spike trains were analyzed using custom-written MATLAB code. Animal tracking position was extracted from video footage by Ethovision XT software (Noldus, Wageningen, the Netherlands) which was synchronized with the electrophysiology data acquisition system. At the end of experiments, electrolytic lesions were made to verify tetrode placement. Brain tissue was fixed by transcardial perfusion and Nissl stained (Figure 1A). Only animals with clear lesions in the CA1 pyramidal layer were included in the analysis.

### Behavioral protocol

An experiment consisted of four sessions (two per day on two consecutive days) during which hippocampal neural ensemble activity was recorded as the mice freely explored a fully transparent, circular open field arena (diameter 64cm) for 30min. The arena was surrounded by black curtains and four large geometrical visual cues (Figure 1B). In the final (fourth) session, three of the visual cues were removed (“Probe” session). The two daily recording sessions were separated by a two-hour break, during which the animal rested in its home cage. Each animal was used for multiple (consecutive) experiments; a new set of visual wall cues was selected for each iteration.

### Neuronal analysis

Periods of inactivity (animal speed <3cm/s) were excluded from analysis. Videotracking data were visually inspected, checked for accuracy, and corrected manually when necessary. Recording stability of individual clusters of spikes was examined; clusters whose first principal component drift exceeded more than three standard deviations across both sessions within a day were excluded from analysis. Classification of putative pyramidal cells was based on their firing rate and16 the mean of the autocorrelogram, as previously described by our lab^32^.

### Place cell analysis

To create firing maps of individual neurons (Figure 1D), spike data were plotted on binned arena occupancy data (pixels: 2x2cm), normalized by the total time spent in each bin, and smoothed (radius: 2). Bins that received insufficient sampling (<200ms) were excluded from analysis. Only neurons that displayed place-related activity in at least one session were included in analysis. Place fields were defined as areas larger than 10 adjacent pixels where a pyramidal cell exhibited more than 30% of its maximum firing rate. Spatial information per spike was calculated as described in ^48^. Spatial specificity was calculated as the firing rate increase of each cell within its field (in-field firing rate divided by out-field firing rate).

## Acknowledgements

We thank L. Noldus for the use of Ethovision XT software, K. Harris for the use of Klustakwik, and L. Hazan for the use of Klusters. Animals were kindly provided by Prof. Dr. R. Willemsen at the Department of Clinical Genetics, Erasmus MC in Rotterdam, The Netherlands. This work was supported by SenterNovem BSIK grant 03053, STW grant AET7613, and EU project 720270 – HBP SGA1 (Human Brain Project) to C.M.A.P.

## Author contributions

T.A., C.M.A.P., and F.P.B. designed the experiments. T.A. performed the experiments. T.A. andF.P.B. analyzed the data and wrote the manuscript.

